# Neural responses prior to licking onset in the striatal matrix compartment in mice

**DOI:** 10.1101/2025.05.23.655747

**Authors:** Yuta Ishimaru, Tomohiko Yoshizawa, Taishi Kimoto, Tadashi Inui, Yasutaka Yawaka, Makoto Funahashi

## Abstract

Licking is a continuous tongue thrust observed during drinking in rodents and humans and is often studied as an essential tongue movement for feeding and swallowing. The striatum, a component of the basal ganglia, plays a critical role in licking onset; however, it is unclear how the two compartments of the striatum—the matrix and striosomes—contribute to the control of licking onset. In this study, we used male and female transgenic mice that selectively expressed Cre recombinase in matrix or striosome neurons and subjected them to operant conditioning based on licking of a spout, during which neuronal activity in both compartments was measured using fiber photometry. Only matrix neurons showed responses prior to licking onset. In addition, the matrix neural response before licking onset was larger when mice licked a spout ipsilateral to the recording hemisphere of the brain than that observed when licking the contralateral spout. This response was observed similarly in mice conditioned to receive a reward regularly and those conditioned to receive a reward randomly, suggesting that the response was unrelated to whether the reward was predictable or unpredictable. Matrix neural activity was negatively correlated with the number of licks during the water intake behavior following the first lick. These findings suggest that matrix neurons are involved in the preparatory process for licking onset as well as in the regulation of licking frequency during water intake.

**Significant Statement:** This study demonstrated that during the expression of operant conditioning behaviors based on licking, striatal matrix neurons showed responses prior to licking onset. Additionally, these responses were larger when the mouse licked the spout ipsilateral to the brain hemisphere undergoing recording than when a spout in the contralateral direction was licked. This result was also true for mice conditioned using either regular or random reward conditions. Additionally, the number of licks during water ingestion behavior following the initial lick was negatively correlated with matrix neuron activity. These changes in matrix neuron activity are suggested to be involved in the preparatory process for licking onset, independent of reward prediction, and in the regulation of licking frequency during drinking.

## Introduction

Tongue movements are integral to the processes of eating and swallowing, particularly for the formation and transportation of the bolus (Hiiemae and Palmer, 1999). An appropriate model for studying tongue movements is licking behavior, which consists of a continuous tongue thrusting motion. Licking behavior is generated by the central pattern generators in the brainstem (Travers et al., 1997), which are regulated by top-down signals from the basal ganglia (Deniau and Chevalier, 1992; Redgrave et al., 1992; Shammah-Lagnado et al., 1992; Rossi et al., 2016; Toda et al., 2017). The striatum is a major cortical input site of the basal ganglia. Previous studies in rodents demonstrated that their licking movements were impaired by dopamine deficiency in the striatum (Skitek et al., 1999; Ciucci et al., 2011; Chen et al., 2019), and that stimulation of direct and indirect pathway striatal neurons initiated and suppressed licking, respectively (Bakhurin et al., 2020). The striatum consists of two neurochemically and anatomically distinct compartments: the matrix, which is rich in calbindin (Dong et al., 2025) and receives inputs from the sensorimotor and associative cortices, and the striosomes (also known as patches), which are rich in µ-opioid receptors and prodynorphin (Cui et al., 2014), receive input from the limbic cortex, and monosynaptically project to midbrain dopaminergic neurons (Gerfen, 1984, 1989; Jiménez-Castellanos and Graybiel, 1989; Eblen and Graybiel, 1995; Kincaid and Wilson, 1996). Although previous *in vivo* calcium imaging studies have shown that striosomal neural activity correlates with the number of licks during reward intake in classical conditioning (Bloem et al., 2017; Yoshizawa et al., 2018), it remains unclear which compartment plays a more dominant role in the control of licking movements.

Therefore, in the present study, we selectively recorded striatal neural activities from the matrix and striosome compartments of mice during left and right licking movements and then analyzed licking-related neural activities. Matrix neurons responded before the onset of licking, and the responses were larger when mice licked the waterspout ipsilateral to the brain hemisphere being recorded than when licking the spout on the contralateral side. This activity was not affected by reward prediction. Our findings suggest that matrix neurons are more dominant than striosomal neurons in controlling tongue movements.

## Materials and Methods

### Animals

The Hokkaido University Animal Use Committee approved this study. Male and female Calb1-IRES-Cre (129S-Calb1tm2.1(cre)Hze/J, The Jackson Laboratory Cat# 028532; four male mice, one female mouse; 8–10 weeks old) and Pdyn-IRES-Cre mice (129S-Pdyn(tm1.1(Cre)/Mjkr)/LowlJ, The Jackson Laboratory Cat# 027958; three male mice, two female mice; 8–10 weeks old) were housed individually under a 12/12 h light/dark cycle (lights on at 7 A.M.; off at 7 P.M.). Experiments were performed during the light phase. Water intake was restricted to 1–2 mL/day for 2 days before and during the experiments. Food was provided *ab libitum* for the entire period.

### Surgery

Mice were anesthetized with isoflurane (1.0%–4.0%) and placed in a stereotaxic frame. The skull was exposed, and a hole was drilled in the skull. For fiber photometry recordings, AAV5.CAG.Flex.GCaMP6f.WPRE.SV40 (left hemisphere: five mice, right hemisphere: five mice, 100835-AAV5, Addgene, Watertown, MA, USA) was injected into the dorsomedial striatum (DMS) (AP: +0.5, ML: 1.75, DV: 2.85 mm from the brain surface, volume: 400 nL) using a microsyringe pump (Legato100, Kd Scientific, Holliston, MA, USA). After adeno-associated virus (AAV) injection, an optical probe (diameter: 400 μm, length: 5.0 mm, R-FOC-BL400C-50NA, RWD, Guangdong, China) was implanted 200 μm above the AAV injection coordinates (AP: +0.5, ML: 1.75, DV: 2.65 mm from the brain surface). The optical fiber was then fixed with adhesive dental cement (Super Bond, Sun Medical, Shiga, Japan). A head plate (CF-10, Narishige, Tokyo, Japan) was fixed with pink dental cement (Unifast 2, GC, Tokyo, Japan). Analgesics and antibiotics were applied postoperatively as required (meloxicam, 1 mg/kg s.c.; 0.1% gentamicin ointment, *ad usum externum*).

### Behavioral tasks

The heads and bodies of mice were restricted using a head plate and a metal tube, respectively, and spouts were placed on both the left and right sides of their mouths (Fig. 1A). Licks were detected by interruptions of an infrared beam placed in front of the water tube. In the ipsilateral block, water was delivered from the spout on the side ipsilateral to the recording hemisphere, whereas in the contralateral block, it was delivered from the contralateral side. Each trial began by lighting a light-emitting diode (LED) (Fig. 1B). When mice spontaneously licked the spout on the appropriate side of the block, a drop of 5% sucrose water (4 µL) was immediately presented. At the end of a trial, the LED was turned off, followed by a 10±3 s inter-trial-interval. A daily session consisted of a 20-min ipsilateral block and a 20-min contralateral block. An additional experiment was performed to measure the effect of reward prediction on neural activity, which consisted of a 40-min session. When mice spontaneously licked the waterspout on the ipsilateral side, a drop of 5% sucrose water was alternately delivered or not delivered after a 0.5 s delay. The other behavioral task components were similar to those performed in the first experiment.

**Figure 1.**
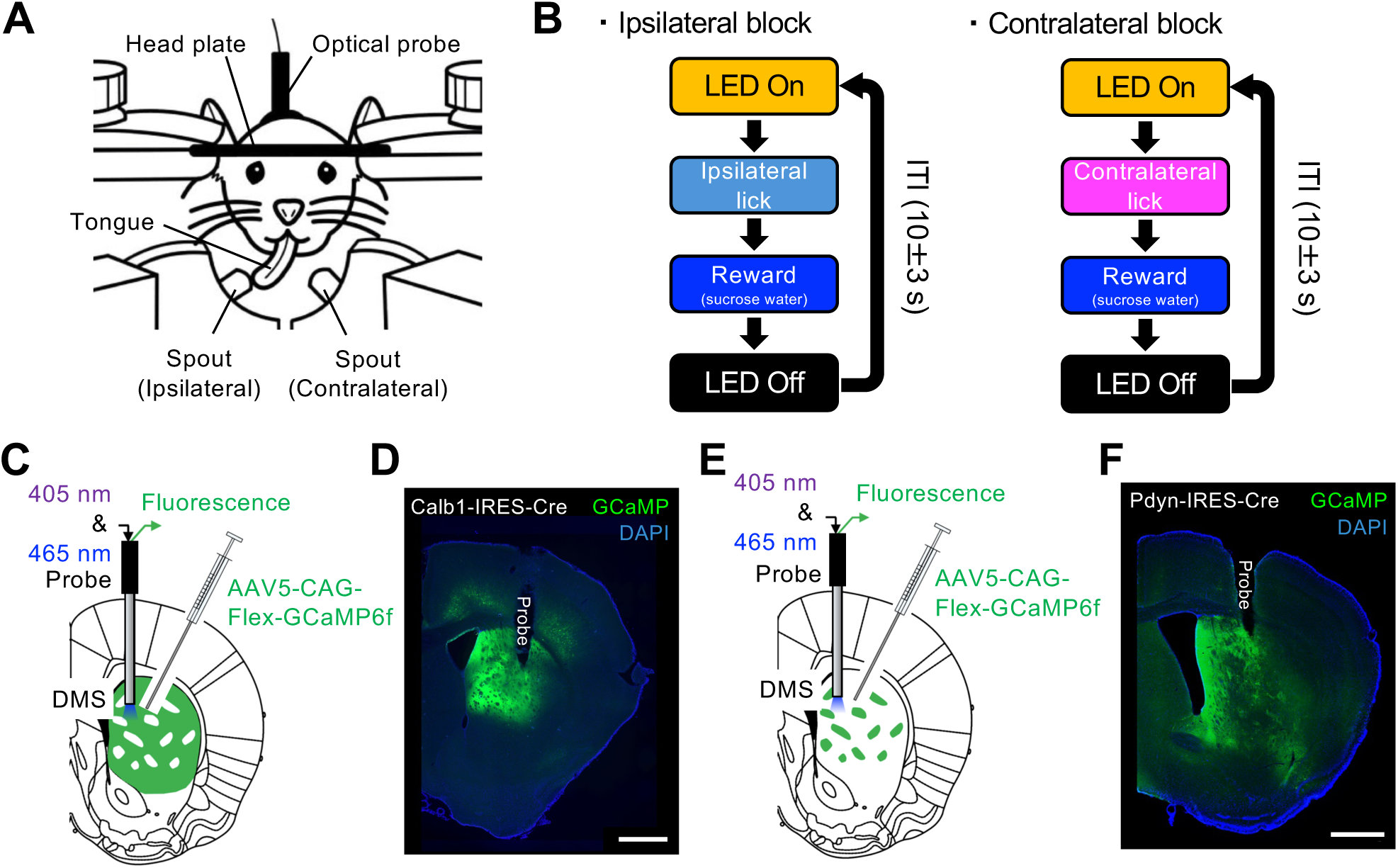
| Measurement of GCaMP fluorescence in the matrix and striosomes during operant conditioning. **A.** Schematic illustration of the behavioral apparatus. The head and body of the mouse were restricted by a metal frame and tube. The two spouts were placed to the left and right of the mouth. Spout-licking behaviors were monitored using an infrared sensor. An optical fiber was connected to the optical probe implanted in the dorsomedial striatum (DMS) for fiber photometry of GCaMP6f. **B.** Diagram of an operant conditioning task. Each trial began when a light-emitting diode (LED) was illuminated. In the ipsilateral and contralateral blocks, a drop of sucrose water was delivered immediately after the mice licked a waterspout placed on the ipsilateral or contralateral side, respectively, of the fiber-implanted hemisphere. After water delivery, the LED was turned off, followed by an inter-trial interval (ITI). **C.** Schematic illustration of the measurement of GCaMP fluorescence from matrix neurons. GCaMP6f was selectively expressed in matrix neurons via injection of AAV5.CAG.Flex.GCaMP6f into the DMS of Calb1-IRES-Cre mice. An optical probe was implanted in the DMS to measure the calcium-dependent fluorescence of GCaMP6f excited by a 465 nm LED. **D.** Histological image of Cre-dependent GCaMP6f-expressing neurons in the striatum of a Calb1-IRES-Cre mouse. Scale bar: 500 µm. **E.** Schematic illustration of the measurement of GCaMP fluorescence from striosome neurons. GCaMP6f was selectively expressed in striosome neurons via injection of AAV5.CAG.Flex.GCaMP6f into the DMS of Pdyn-IRES-Cre mice. **F.** Histological image of Cre-dependent GCaMP6f-expressing neurons in the striatum of a Pdyn-IRES-Cre mouse. GCaMP6f was mosaically expressed in the striatum. Scale bar: 500 µm.

### Fiber photometry

The fiber photometry system consisted of 465 and 405 nm excitation channels that were used to obtain a calcium-dependent signal and a calcium-independent isosbestic signal, respectively. Fluorescence from GCaMP6f and isosbestic fluorescence were directed with dichroic mirrors (iFMC6_IE(400-410)_E1(460-490)_F1(500-540)_E2(555-570)_F2*(580-680)_S, Doric) and were acquired using a photodetector. The signals were passed through a 10× amplifier and were sampled at 1 kHz using a data acquisition system (Power1401, Cambridge Electronic Design, Cambridge, UK). The acquired photometry signals were processed using custom-written MATLAB code (MATLAB R2018a, MathWorks, Natick, MA, USA). The detailed protocol was described in our recent paper (Yoshizawa and Funahashi, 2025).

### Immunohistochemistry

After all experiments were completed, the mice were deeply anesthetized with pentobarbital sodium and then perfused with 4% paraformaldehyde. Brains were carefully removed so that the optical fibers would not cause tissue damage, post-fixed in 4% paraformaldehyde at 4°C overnight, and then transferred to a 30% sucrose/0.1M phosphate buffer solution at 4°C until the brains sank to the bottom. Coronal sections including the striatum were cut at a thickness of 50 μm on a freezing microtome (REM-710; Yamato, Saitama, Japan). Free-floating sections were washed four times in phosphate-buffered saline (PBS) for 15 min and placed in blocking buffer containing 10% normal donkey serum (017-000-121, Jackson ImmunoResearch Laboratories, West Grove, PA, USA) and 0.1% Triton X-100 in PBS for 1 h at room temperature. The sections were simultaneously incubated in chicken anti-GFP primary antibody (GFP-1010, Aves Labs, Davis, CA, USA) diluted 1:500 in blocking buffer overnight at 4°C. Afterward, sections were washed four times for 15 min in PBS. The sections were then incubated in donkey anti-chicken Alexa Fluor 488 secondary antibody (703-545-155, Jackson ImmunoResearch Laboratories) diluted 1:500 in blocking buffer for 2 h at room temperature. The sections were washed four times for 15 min in PBS, mounted on glass slides, and coverslipped with VECTASHIELD Mounting Medium with DAPI (Vector Laboratories, Newark, CA, USA). A fluorescence microscope (Eclipse Ci-L, Nikon, Tokyo, Japan) was used to inspect the stained tissue, and images were obtained using NIS-Elements software (NIS-Elements D, Nikon).

### Experimental design and statistical analysis

The analyses include 7164 behavioral and neural trials recorded over a total of 57 sessions with 10 mice. We used appropriate statistical tests when applicable, i.e., paired or unpaired t-tests and Pearson correlation analysis with or without Bonferroni’s multiple comparisons tests. Differences were considered statistically significant when p<0.05. Details are described in the Results section.

## Results

### Licking-related neural activities in the matrix compartment during operant conditioning

Head-fixed mice performed an operant conditioning task (Fig. 1A, B). Transgenic mice (Calb1-IRES-Cre) selectively expressing Cre in their matrix neurons (Evans et al., 2020) were employed to record licking-related neural activity from the matrix. After Cre-dependent expression of the genetic calcium sensor GCaMP6f (Chen et al., 2013) was induced by AAV injection into the DMS (Fig. 1C), fiber photometry recordings were performed during the task. In all five Calb1-IRES-Cre mice, we confirmed that GCaMP6f-expressing neurons were located at the tip of the optical fiber (Fig. 1D). Figure 2(A, B) shows representative licking behavior and GCaMP fluorescence recorded from the matrix of the left hemisphere. When the mouse licked the ipsilateral (left) and contralateral (right) spouts, the fluorescence increased before the onset of the first lick after the LED was illuminated. This increased fluorescence continued during the water ingestion licks. The average fluorescence during the pre-licking period (−1.0 to 0 s before the onset of the first lick) was significantly larger than that during the baseline period (−2.0 to −1.0 s before the onset of the first lick) in both blocks (ipsilateral: −0.46±0.054, baseline and 0.044±0.086, pre-licking, p=1.2e−07; contralateral: −0.39±0.061, baseline and 0.28±0.10, pre-licking, p=1.6e−07, paired t-test, all fluorescence was measured using z-scores, Fig. 2C). There was no significant difference between the ipsilateral and contralateral blocks regarding the average fluorescence during the pre-licking and licking (0 to 1.5 s after the onset of the first lick) periods (pre-licking: 0.044±0.086, ipsilateral and 0.28±0.10, contralateral, p=0.084; licking: 0.52±0.045, ipsilateral and 0.38±0.081, contralateral, p=0.12, unpaired t-test, Fig. 2D, E). In the ipsilateral block, there was no significant correlation between the number of licks and the average fluorescence during the licking period (r=−0.17, p=0.11, Pearson correlation analysis, Fig. 2F), whereas in the contralateral block, there was a significant negative correlation (r=−0.52, p=1.2e−05).

**Figure 2.**
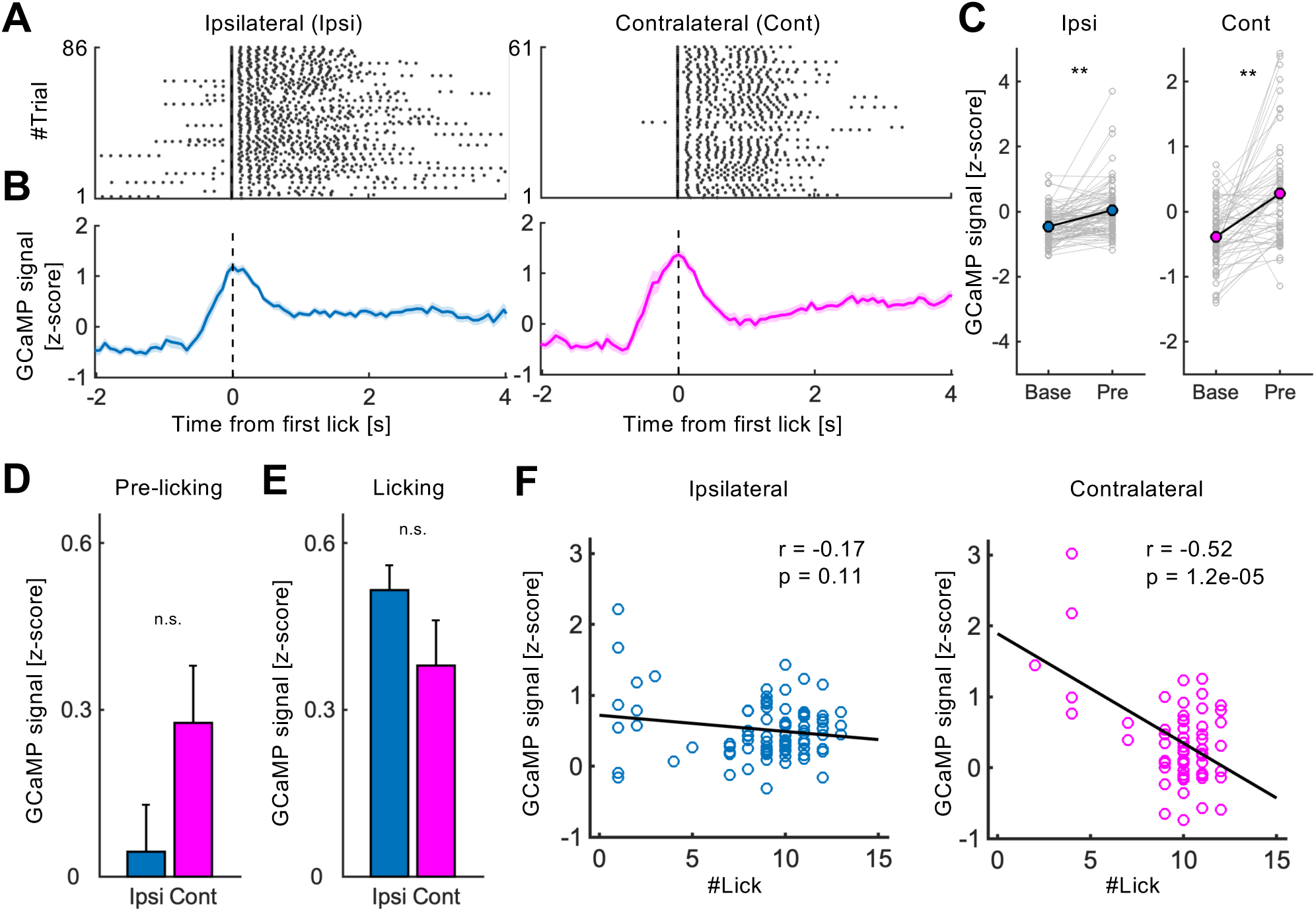
| Representative activity of matrix neurons during operant conditioning. **A.** Spout-licking behavior in the ipsilateral and contralateral blocks, sorted by the first lick after LED illumination. Black dots indicate the timing of spout-licking behaviors. **B.** Average GCaMP fluorescence recorded from matrix neurons in the ipsilateral and contralateral blocks. The fluorescence was recorded in the same session as the events in **A.** Matrix neurons showed responses prior to the onset of the first lick in both blocks. **C.** Average GCaMP fluorescence during the baseline (Base) and pre-licking (Pre) periods. **: p<0.01, paired t-test. **D.** Comparison of average GCaMP fluorescence during the pre-licking period between the ipsilateral and contralateral blocks. Error bars indicate the SEM. n.s.: p>0.05, unpaired t-test. **E.** Comparison of average GCaMP fluorescence during the licking period between the ipsilateral and contralateral blocks. n.s.: p>0.05, unpaired t-test. **F.** Correlation between the number of licks and average GCaMP fluorescence during the licking period. The average fluorescence during the licking period is plotted against the number of licks in the same period. Circles and the black line indicate the average GCaMP fluorescence in each trial and the regression line, respectively. Pearson correlation analysis.

### Licking-related neural activity in the striosome compartment during operant conditioning

Transgenic mice (Pdyn-IRES-Cre) selectively expressing Cre in their striosomal neurons (Evans et al., 2020; Xiao et al., 2020; Yoshizawa and Funahashi, 2025) were employed to record licking-related neural activity from the striosomes. After Cre-dependent expression of GCaMP6f was induced by AAV injection into the DMS, fiber photometry recordings were performed during the task (Fig. 1E). We confirmed that GCaMP6f-expressing neurons were located at the tip of the optical fiber in all five Pdyn-IRES-Cre mice (Fig. 1F). Figure 3(A, B) shows representative licking behavior and GCaMP fluorescence recorded from striosomes in the right hemisphere. In contrast to the matrix, increased fluorescence during the pre-licking period was not observed in either the ipsilateral or contralateral blocks (ipsilateral: −0.067±0.085, baseline and −0.059±0.093, pre-licking, p=0.84; contralateral: −0.061±0.047, baseline and −0.027±0.047, pre-licking, p=0.53, paired t-test, Fig. 3C). The fluorescence peaked during the licking period. There was no significant difference between the ipsilateral and contralateral blocks regarding the average fluorescence during the licking period (ipsilateral: 0.69±0.072, contralateral: 0.80±0.053, p=0.23, unpaired t-test, Fig. 3D). In both the ipsilateral and contralateral blocks, a significant positive correlation was observed between the number of licks and the average fluorescence during the licking period (ipsilateral: r=0.42, p=0.0032; contralateral: r=0.39, p=6.2e-05, Pearson correlation analysis, Fig. 3E).

**Figure 3.**
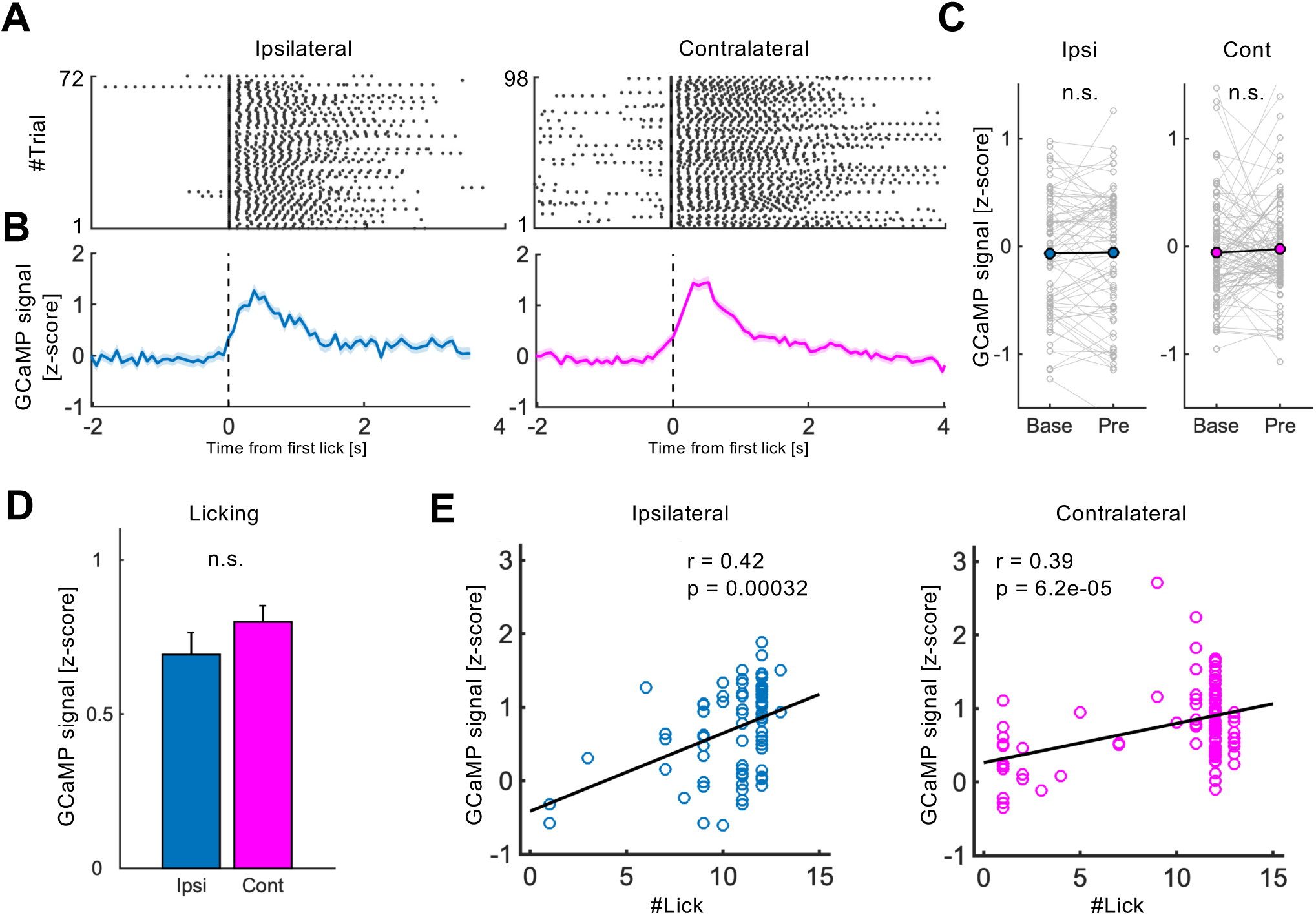
| Representative activity of striosome neurons during operant conditioning. **A.** Spout-licking behavior in the ipsilateral and contralateral blocks, sorted by the first lick after LED illumination. Black dots indicate the timing of spout-licking behaviors. **B.** Average GCaMP fluorescence recorded from striosome neurons in the ipsilateral and contralateral blocks. The fluorescence was recorded in the same session as the events in **A.** Striosome neurons did not show responses prior to the onset of the first lick in either block. **C.** Average GCaMP fluorescence during the baseline (Base) and pre-licking (Pre) period. n.s.: p≥0.05, paired t-test. **D.** Comparison of average GCaMP fluorescence during the licking period between the ipsilateral and contralateral blocks. n.s.: p>0.05, unpaired t-test. **E.** Correlation between the number of licks and average GCaMP fluorescence during the licking period. The average fluorescence during the licking period is plotted against the number of licks during the same period. Circles and the black line indicate the average GCaMP fluorescence in each trial and the regression line, respectively. Pearson correlation analysis.

### Comparison of licking-related neural activities between the matrix and striosome compartments

To quantitatively examine differences in licking-related neural activities between the matrix and striosomes, we first averaged the GCaMP fluorescence of all five Calb1-IRES-Cre mice (Fig. 4A). The average fluorescence during the pre-licking period was significantly greater than that during the baseline period in both the ipsilateral and contralateral blocks (ipsilateral: −0.36±0.017, baseline and 0.30±0.026, pre-licking, p=7.8e−126, n=1318 trials; contralateral: −0.31±0.018, baseline and 0.058±0.021, pre-licking, p=5.0e−83, n=1401 trials, paired t-test, Fig. 4B) and was significantly larger in the ipsilateral block than in the contralateral block (ipsilateral: 0.30±0.026, contralateral: 0.058±0.021, p=9.6e−13, unpaired t-test, Fig. 4C). The fluorescence during the licking period was also significantly larger in the ipsilateral block than in the contralateral block (ipsilateral: 0.69±0.018, contralateral: 0.46±0.019, p=1.9e−17, unpaired t-test, Fig. 4D). Although the correlation coefficient between the number of licks and the average fluorescence during the licking period was not different between the ipsilateral and contralateral blocks (ipsilateral: r=−0.070±0.060, contralateral: r=−0.14±0.064, p=0.49, n=20 sessions, paired t-test, Fig. 4E), the correlation coefficient was significantly negative in the contralateral block (p=0.043, paired t-test).

**Figure 4.**
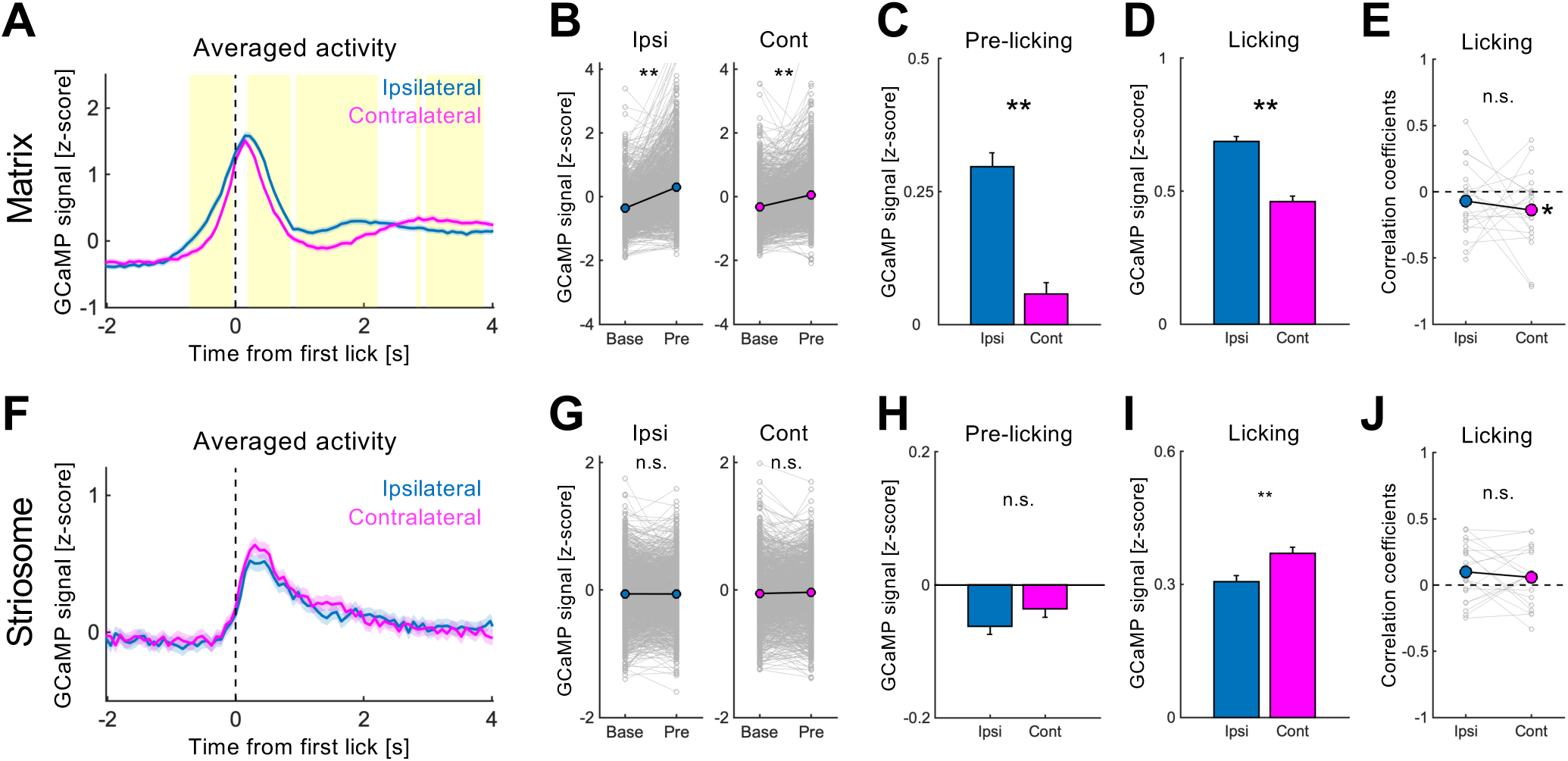
| Population analysis of licking-related neural activity in the matrix and striosomes. **A.** Matrix neural activity averaged over five mice performing an operant conditioning task. GCaMP signals were aligned to the onset of the first lick after LED illumination. Indigo and magenta lines indicate average activity in the ipsilateral and contralateral blocks, respectively. Shaded areas indicate 95% confidence intervals. Yellow bins indicate a significant difference in the z-score between blocks (p<0.01, unpaired t-test followed by Bonferroni correction). **B.** Comparison of matrix neural activity between the baseline (Base) and pre-licking (Pre) periods. Indigo and magenta circles indicate the means of GCaMP signals in the ipsilateral and contralateral blocks, respectively. **: p<0.01, paired t-test. **C, D.** Comparison of matrix neural activity in the pre-licking (C) and licking (D) periods between the ipsilateral and contralateral blocks. Mean ± SEM. **: p<0.01, unpaired t-test. **E.** Correlation coefficients between the number of licks and average GCaMP fluorescence of each mouse from which matrix recordings were performed. Indigo and magenta circles indicate average correlation coefficients of matrix recordings from mice in the ipsilateral and contralateral blocks, respectively. *: p<0.05, n.s.: p≥0.05, paired t-test. **F.** Striosome neural activity averaged over five mice performing the operant conditioning task. **G.** Comparison of striosome neural activity between the baseline (Base) and pre-licking (Pre) periods. n.s.: p≥0.05, paired t-test. **H, I.** Comparison of striosome neural activity during the pre-licking (H) and licking (I) periods between the ipsilateral and contralateral blocks. Mean ± SEM. **: p<0.01, n.s.: p≥0.05, unpaired t-test. **J.** Correlation coefficients between the number of licks and the average GCaMP fluorescence in the striosomes of each mouse. Indigo and magenta circles indicate the average correlation coefficients of mice in which striosomes were recorded in the ipsilateral and contralateral blocks, respectively. n.s.: p≥0.05, paired t-test.

Next, we averaged the GCaMP fluorescence of all five Pdyn-IRES-Cre mice (Fig. 4F). In both the ipsilateral and contralateral blocks, the average fluorescence was not significantly different between the baseline and pre-licking period (ipsilateral: −0.061±0.013, baseline and −0.063±0.012, pre-licking, p=0.85, n=1289 trials; contralateral: −0.0555±0.013, baseline and −0.036±0.013, pre-licking, p=0.12, n=1222 trials, paired t-test, Fig. 4G). There was no significant difference in the average fluorescence of the pre-licking period between the blocks (ipsilateral: −0.063±0.012, contralateral: −0.036±0.013, p=0.14, unpaired t-test, Fig. 4H), whereas during the licking period, the fluorescence was significantly larger in the contralateral block than in the ipsilateral block (ipsilateral: 0.31±0.014, contralateral: 0.37±0.014, p=0.0017, Fig. 4I). The correlation coefficient between the number of licks and the average fluorescence during the licking period was not significantly different between the blocks (ipsilateral: r=0.10±0.049, contralateral: r=0.58±0.048, p=0.47, n=20 sessions, paired t-test, Fig. 4J). The correlation coefficients were not significantly different from zero in either block (ipsilateral: p=0.056, contralateral: p=0.24, paired t-test).

### Effects of reward prediction on the licking-related activity of matrix neurons

The striatum plays a critical role not only in motor control but also in reward prediction (Samejima et al., 2005; Ito and Doya, 2009; Kim et al., 2009; Ito and Doya, 2015; Yoshizawa et al., 2018, 2023). To clarify whether matrix neuron responses prior to licking onset reflected motor-related or reward-predictive neural activity, we recorded matrix neural activity during another operant conditioning experiment in which mice alternately received a reward and no reward after licking the spout on the ipsilateral side (Fig. 5A). This side was used because the pre-licking response was larger in the ipsilateral block than in the contralateral block. In addition, a 0.5 s delay was inserted before the reward presentation to test whether the reward presentation influenced neural activity during the licking period.

**Figure 5.**
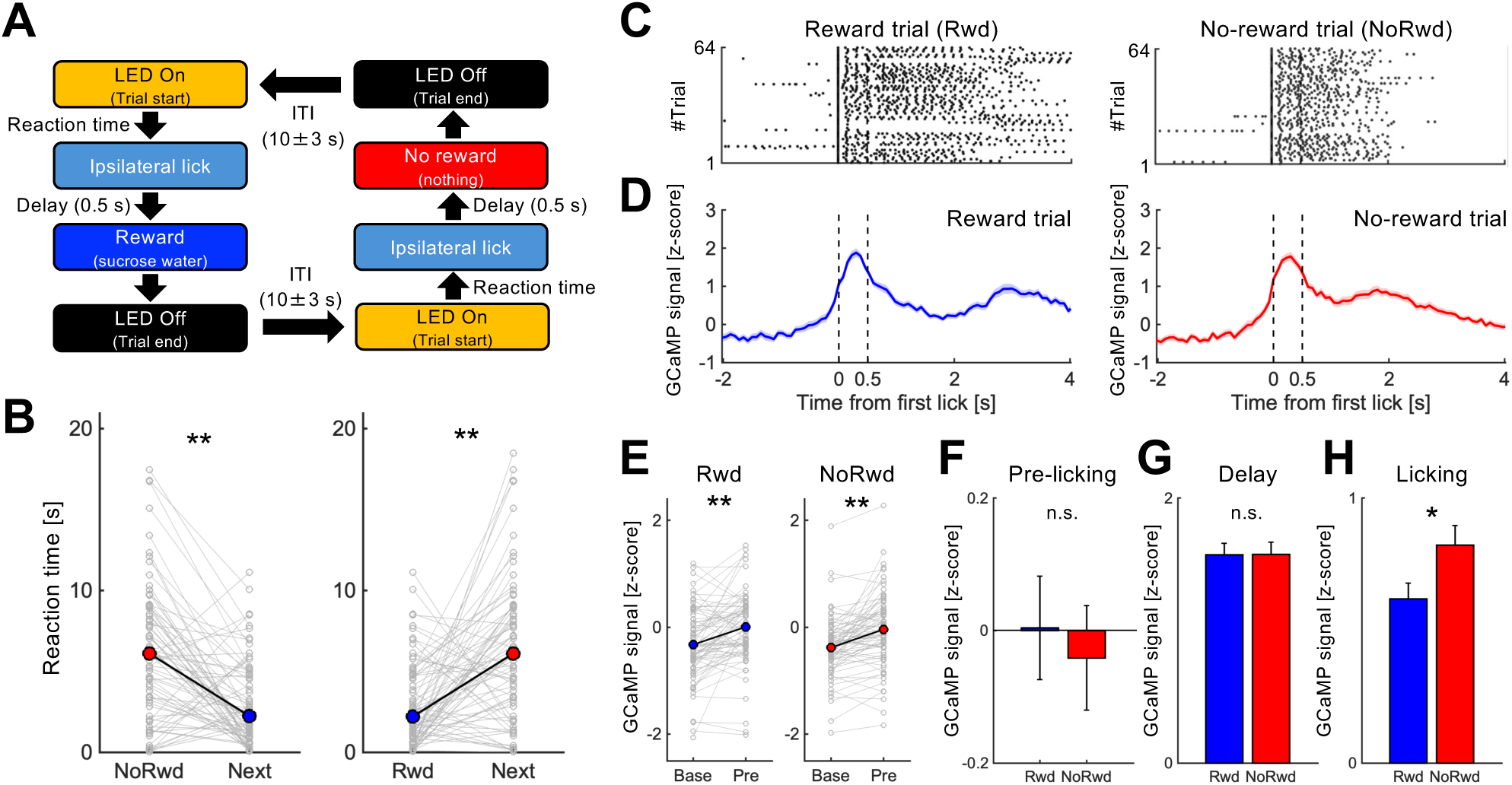
| Effects of reward prediction on licking-related neural activity in the matrix. **A.** Diagram of an operant conditioning task designed to test effects of reward prediction on licking-related neural activity in the matrix. Each trial began with LED illumination. When the mouse spontaneously licked the waterspout placed on the ipsilateral side of the optical probe-implanted hemisphere, a drop of sucrose water was alternately delivered or not delivered after a 0.5 s delay period. At the end of each trial, the LED was turned off, followed by a 10±3 s ITI. **B.** Representative example of the reaction time (RT) in a single session. RTs were shorter in the trials following no-reward (NoRwd) trials, whereas they were longer in the trials following reward (Rwd) trials. Red and blue circles indicate the mean RTs in NoRwd and Rwd trials, respectively. **: p<0.01, paired t-test. **C.** Spout-licking behavior in reward and no-reward trials, sorted by the first lick after LED illumination. Black dots indicate the timing of spout-licking behaviors. **D.** Average GCaMP fluorescence recorded from matrix neurons in reward and no-reward trials. The fluorescence was recorded in the same session as the events in **C**. Matrix neurons showed responses prior to the onset of the first lick in both reward and no-reward trials. Blue and red lines indicate averaged fluorescence in reward and no-reward trials, respectively. Shaded areas indicate 95% confidence intervals. **E.** Average GCaMP fluorescence during the baseline (Base) and pre-licking (Pre) periods. Blue and red circles indicate the mean GCaMP signal in reward and no-reward trials, respectively. **: p<0.01, paired t-test. **F–H.** Comparison of matrix neural activity during the pre-licking (F), delay (G), and licking (H) periods between reward and no-reward trials. Mean ± SEM. **: p<0.01, n.s.: p≥0.05, unpaired t-test.

In each trial, we measured the reaction time (RT) from when the LED was illuminated to the onset of the first lick because under the reward alternation scheme, the RT in the reward trial became shorter than that in the no-reward trial when subjects successfully predicted the upcoming reward (Isomura et al., 2013). The RT was shorter or longer following subsequent no-reward or reward trials, respectively (post-no-reward trials: p=1.5e−05, post-reward trials: p=0.00054, paired t-test, Fig. 5B). Figure 5(C, D) shows representative licking behaviors and GCaMP fluorescence recorded from neurons in the matrix compartment, respectively. The fluorescence was significantly increased during the pre-licking period compared with that during the baseline period in both reward and no-reward trials (reward trial: −0.032±0.076, baseline and 0.0042±0.078, pre-licking, p=6.9e−06; no-reward trial: −0.38±0.065, baseline and −0.041±0.079, pre-licking, p=4.9e−08, paired t-test, Fig. 5E). The average fluorescence was not significantly different between reward and no-reward trials during the pre-licking and delay periods (pre-licking: 0.0042±0.078, reward trial and −0.041±0.079, no-reward trial, p=0.68; delay: 1.6±0.085, reward trial and 1.6±0.093, no-reward trial, p=0.99, unpaired t-test, Fig. 5F, G). During the licking period, the fluorescence was significantly larger in the reward trial than in the no-reward trial (0.62±0.061, reward trial and 0.82±0.075, no-reward trial, p=0.036, Fig. 5H).

To quantitatively examine the difference in the GCaMP signal between reward and no-reward trials, we averaged the trial-by-trial signal of the sessions in which mice successfully predicted the reward and no-reward trials (17 sessions including five mice, Fig. 6A). In both reward and no-reward trials, the average fluorescence was significantly larger during the pre-licking period than during the baseline period (reward trial: −0.15±0.022, baseline and 0.097±0.022, pre-licking, p=1.3e−34, n=966 trials; no-reward trial: −0.32±0.021, baseline and 0.10±0.025, pre-licking, p=5.7e−77, n=968 trials, paired t-test, Fig. 6B). During the pre-licking and delay periods, the average fluorescence was not significantly different between the reward and no-reward trials (pre-licking: 0.097±0.022, reward trial and 0.10±0.025, no-reward trial, p=0.82; delay: 1.08±0.027, reward trial and 1.2±0.028, no-reward trial, p=0.053, unpaired t-test, Fig. 6C, D). During the licking period, the average fluorescence was significantly greater in the no-reward trial than in the reward trial (0.37±0.021, reward trial and 0.50±0.023, no-reward trial, p=2.5e−05, Fig. 6E). These results indicate that increased fluorescence in the pre-licking period reflected motor-related neural activity.

**Figure 6.**
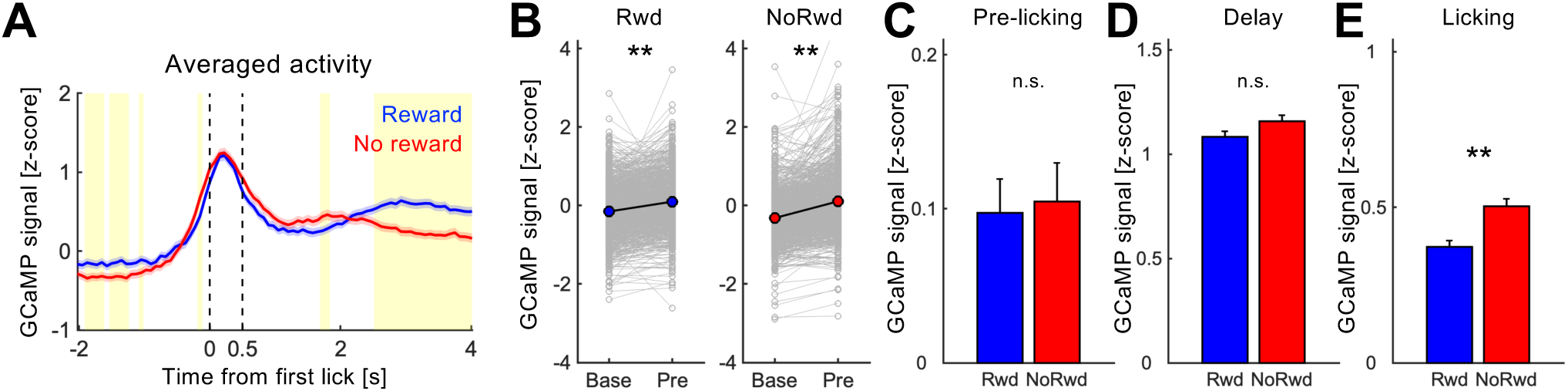
| Population analysis of the effects of reward prediction on licking-related neural activity in the matrix. **A.** Matrix neural activity averaged over five mice performing the reward alternation operant conditioning task. GCaMP signals were aligned to the onset of the first lick after LED illumination. Blue and red lines indicate averaged activity in reward and no-reward trials, respectively. Shaded areas indicate 95% confidence intervals. Yellow bins indicate a significant difference in the z-score between blocks (p<0.01, unpaired t-test followed by Bonferroni correction). **B.** Comparison of matrix neural activity between the baseline (Base) and pre-licking (Pre) periods. Blue and red circles indicate the mean GCaMP signals in reward and no-reward trials, respectively. **: p<0.01, paired t-test. **C–E.** Comparison of matrix neural activity during the pre-licking (C), delay (D), and licking (E) periods between reward and no-reward trials. Mean ± SEM. **: p<0.01, n.s.: p≥0.05, unpaired t-test.

## Discussion

In the present study, we compared licking-related neural activity between the matrix and the striosome compartments. The major findings are as follows: (1) a neural response prior to licking onset was observed in the matrix but not in the striosomes; (2) this pre-licking neural response was larger when mice licked the spout on the side ipsilateral to the recording hemisphere than that observed when mice licked a spout on the contralateral side; (3) in the matrix, the neural response during the water ingestion licks was negatively correlated with the number of licks, while in the striosomes, no correlation was observed; and (4) the pre-licking neural response in the matrix did not reflect reward prediction.

Matrix neurons have been hypothesized to play a more dominant role in motor control than striosome neurons because the matrix receives input from the motor cortices (Graybiel and Matsushima, 2023). In support of this hypothesis, chemogenetic inactivation of matrix neurons impaired the performance of a learned reach-to-grasp ability (Lopez-Huerta et al., 2016). A more recent study showed that matrix neurons exhibit early activation at the onset of locomotion and that optogenetic activation of matrix neurons promotes locomotion (Dong et al., 2025). In the present study, an increased neural response prior to licking onset was only observed in the matrix, indicating the importance of the matrix in motor control of the tongue.

Electrophysiological studies reported that pyramidal neurons in the primary motor cortex of monkeys (Tanji and Kurata, 1982) and medium spiny neurons in the dorsolateral striatum (DLS) of rats (Isomura et al., 2013) showed firing activity 0.5 to 1.0 s before the onset of hand movements. Such firing is thought to be a preparatory activity for movement initiation. The increased GCaMP fluorescence in the present study also occurred from approximately 1.0 s before licking onset; therefore, there is a possibility that the increase reflected firing activity in preparation for movement initiation.

Licking behavior is a continuous tongue thrusting motion caused by contraction of the genioglossus muscle (Travers and Jackson, 1992). The genioglossus muscle comprises a matched pair of extrinsic muscles of the tongue, originates from the mental spine of the mandible, and terminates within the tongue. Bilateral contraction of the genioglossus muscle causes forward protrusion of the tongue, whereas unilateral contraction causes protrusion to the contralateral side (McClung and Goldberg, 2000; Pittman and Bailey, 2009). The hypoglossal nerve innervates the ipsilateral genioglossus muscle and receives bilateral supranuclear inputs (Ugolini, 1995; Zhang et al., 2014). Therefore, it seems paradoxical that patients with unilateral stroke describe tongue deviation to the healthy side (Umapathi et al., 2000; Wei et al., 2012). To explain this phenomenon, previous studies have pointed to a bilateral asymmetry in the supranuclear innervation of the hypoglossal nucleus (Lin and Barkhaus, 2009; Morecraft et al., 2014). To our knowledge, the present study is the first report to demonstrate the asymmetry of licking-related activity in the matrix. This result supports the bilateral asymmetry in the supranuclear innervation of the hypoglossal nucleus.

The activity of matrix neurons during water ingestion licks was negatively correlated with the number of licks (Fig. 2F, 4E) and was greater in the no-reward trial than in the reward trial (Fig. 5H, 6E). The licking duration was also shorter in the no-reward trial than in the reward trial (Fig. 5C). These results suggest that activation of matrix neurons during water ingestion licks inhibits licking behavior. In contrast to the matrix, striosome neuron activity during the water ingestion licks was not correlated with the number of licks (Fig. 4J). A recent study reported that chemogenetic stimulation of striosomal neurons inhibited contralateral rotation and the total distance traveled in the task (Okunomiya et al., 2025). Therefore, striosomal neurons might play a different role in licking and locomotion.

Many studies have demonstrated that striatal neurons contribute to the prediction of future reward. For instance, electrophysiological and *in vivo* calcium imaging studies have shown that value information of reinforcement learning is represented in neural activity in the striatum of monkeys and rodents (Samejima et al., 2005; Ito and Doya, 2009, 2015; Yoshizawa et al., 2018). To clarify whether the matrix neuron response prior to licking onset was motor-related activity or reward-predictive activity, we recorded matrix neural activity under the reward alternation paradigm, in which mice can easily predict a reward or no reward (Isomura et al., 2013). Their reaction time after turning on an LED was shorter in the reward trial than in the no-reward trial because they wanted to receive a reward as soon as possible. This result indicates that the mice were able to predict the upcoming reward. The pre-licking neural response in the matrix, however, was not significantly different between the reward and no-reward trials, suggesting that the pre-licking response did not reflect reward prediction, but rather preparation for licking onset. Moreover, DLS firing activity prior to onset of hand movement has been reported to be modulated by reward prediction (Isomura et al., 2013). The licking-related neural activity of the DLS matrix might be modulated by reward prediction.

## Acknowledgements

This work was supported by JSPS KAKENHI Grant Numbers JP22K15247 and JP25K09843, the Northtec Foundation, and the generous research support of Hokkaido University for young researchers. We thank Edanz (https://jp.edanz.com/ac) for editing the English text of a draft of this manuscript.

